# The global rarity of intact coastal regions

**DOI:** 10.1101/2021.05.10.443490

**Authors:** Brooke A Williams, James E M Watson, Hawthorne L Beyer, Carissa J Klein, Jamie Montgomery, Rebecca K Runting, Leslie A Roberson, Benjamin S Halpern, Hedley S Grantham, Caitlin D. Kuempel, Melanie Frazier, Oscar Venter, Amelia Wenger

## Abstract

Management of the land-sea interface is considered essential for global conservation and sustainability objectives, as coastal regions maintain natural processes that support biodiversity and the livelihood of billions of people. However, assessments of coastal regions have focused on either strictly the terrestrial or marine realm, and as a consequence, we still have a poor understanding of the overall state of Earth’s coastal regions. Here, by integrating the terrestrial human footprint and marine cumulative human impact maps, we provide a global assessment of the anthropogenic pressures affecting coastal areas. Just 15.5% of coastal areas globally can be considered having low anthropogenic pressure, mostly found in Canada, Russia, and Greenland. Conversely, 47.9% of coastal regions are heavily impacted by humanity with most countries (84.1%) having >50% of their coastal regions degraded. Nearly half (43.3%) of protected areas across coastal regions are exposed to high human pressures. In order to meet global sustainability objectives, we identify those nations that must undertake greater actions to preserve and restore coastal regions so as to ensure global sustainable development objectives can be met.

## Introduction

Coastal regions encompass some of the most biodiverse and unique ecosystems on Earth, including coral reefs, kelp forests, seagrass, tidal flats, mangroves, estuaries, salt marshes, wetlands, and coastal wooded habitat (Ray 1991). The persistence of many species relies upon processes that occur across the coastal region including breeding, foraging and migration of both terrestrial and aquatic species (and species that inhabit both systems), nutrient exchange, riverine inputs, and tidal flow (Hazlitt et al. 2010; Fang et al. 2018). Coastal ecological processes underpin critical ecosystem services to humanity like fisheries (Barbier et al. 2011), storm protection (Barbier 2015), and carbon storage and sequestration (known as “blue carbon”) to help mitigate climate change (Mcleod et al. 2011). As a consequence, intact coastal regions (i.e., those that have relatively low human pressure) are critical for maintaining natural processes that support biodiversity and ecosystem services (Fang et al. 2018). These ecosystem services are relied upon by billions of people for their livelihoods (Cinner 2014) and wellbeing (Vo et al. 2012).

Coastal degradation from anthropogenic activity has resulted in profound declines in biodiversity and ecosystem services. For example, the destruction of coastal habitats is leading to declines in adult coral reef fish populations, such as the economically valuable bumphead parrotfish (*Bolbometopon muricatum*; Hamilton et al. 2017). Many species can only persist in coastal ecosystems with high levels of ecological integrity. The marbled murrelet (*Brachyramphus marmoratus*), a seabird from the North Pacific, is an example of a species that relies upon old growth conifer forests to nest (up to 30 km from the shoreline) and high-quality marine habitats to forage (mainly within 500 m of the shoreline; Hazlitt et al. 2010). Ecosystem services are also being lost. It is estimated that coastal degradation leads to 0.15 - 1.02 billion y^-1^ tonnes of carbon dioxide being released from coastal ecosystems and direct economic damages arising from the loss of vegetated coastal ecosystems estimated at $6 - 42 billion USD y^-1^ (Pendleton et al. 2012).

With as much as 74% of the world’s population living within 50 km of the coast (Small & Nicholls 2003), understanding spatial patterns of human influence on coastal regions is essential for identifying natural ecosystems that may be in crisis and require conservation action, to ensure the long-term persistence of important ecological coastal processes (Halpern et al. 2015). Human pressure maps have been developed for the terrestrial and marine realms to inform conservation and management of biodiversity and ecosystem services. At a global scale, the terrestrial human footprint (Venter et al. 2016b; Williams et al. 2020) and the marine cumulative human impact maps (Halpern et al. 2019) are a collation of information from eight (e.g., built human environments, population density) and fourteen large scale pressures (e.g., fishing activities, nutrient pollution), respectively. These maps have been used to identify areas with low human pressure at different times, but these analyses have been carried out separately for the land (Allan et al. 2017) and sea (Jones et al. 2018). Consequently, despite significant increases in spatial information on humanity’s impact across Earth, we still have a poor understanding of the state of coastal regions globally.

Quantifying the loss of ecosystem condition and function is challenging. In lieu of formal scientific surveys, proxy indicators can inform initial assessments for planning purposes (Watson & Venter 2019). Here, we develop a coastal region intactness metric that is derived from an integration of terrestrial and marine intactness metrics. We combine the latest terrestrial and marine cumulative pressure maps for the year 2013 to provide the first global assessment of human pressures on Earth’s coastal regions, whereby a coastal region is defined as the transition between marine and terrestrial environments mapped at a 1 km resolution, up to 50 km on either side of the shoreline (Fang et al. 2018). We assess coastal region intactness at the global and national scales, ascertaining which nations contain Earth’s remaining intact coastal regions, and which have the greatest amounts that are degraded. We then quantify human pressure across 11 coastal ecosystems to identify those most at risk, and assess current levels of coastal protection. A Post-2020 Global Biodiversity Framework will be soon agreed at the fifteenth conference of the parties to the Convention on Biological Diversity with the goal of preventing the catastrophic loss of global biodiversity that delivers multiple benefits to humanity (Díaz et al. 2019; Maxwell et al. 2020). Maintaining and restoring coastal ecological integrity is key to meeting this goal, as well as other global sustainability goals outlined by the United Nations’ Sustainable Development Goals (Neumann et al. 2017). Therefore, it is the right moment to evaluate the intensity of human pressure along coastal regions, and to contextualise this assessment against the background of global sustainability objectives to prioritise actions nations can undertake towards retaining, sustainably managing, and restoring coastal regions to the benefit of both biodiversity and the people who rely on them for survival.

## Methods

### Human pressure across coastal regions

To represent human pressure on terrestrial Earth we used the recently released human footprint for the year 2013, which includes pressures on (1) the extent of built human environments, (2) population density, (3) electric infrastructure, (4) crop lands, (5) pasture lands, (6) roadways, (7) railways, and (8) navigable waterways (Venter et al. 2016a; Williams et al. 2020). For the marine environment, we used the cumulative human index, also for the year 2013 which includes pressures from four primary categories, (1) fishing: commercial demersal destructive, commercial demersal non-destructive high bycatch, commercial demersal non-destructive low bycatch, pelagic high bycatch, pelagic low bycatch, artisanal, (2) climate change: sea surface temperature, ocean acidification, sea-level rise (though this component was only used in the sensitivity analysis), (3) ocean: shipping, and (4) land-based: nutrient pollution, organic chemical pollution, direct human (population density), light (Halpern et al. 2019).

For the terrestrial human footprint (1 km^2^ resolution), as per previous studies we defined intact land as anything below a threshold of <4 (Beyer et al. 2019; Williams et al. 2020). This threshold has been found to be robust from a species conservation perspective because, once surpassed, species extinction risk increases dramatically (Di Marco et al. 2018), and several ecosystem processes are altered (Crooks et al. 2017; Di Marco et al. 2018; Tucker et al. 2018). To identify intact areas within the marine realm we regard any value below the 40% quantile which equates to a threshold of 3.87e-2 (of a total range of 0-12; Halpern et al. 2019), for the global cumulative human impact map (1 km^2^ resolution). Following Jones et al. 2018 (Jones et al. 2018) we excluded climate change variables (temperature and UV anomalies, ocean acidification, and sea-level rise) from the marine cumulative human pressure dataset, because the impacts of climate change are widespread and unmanageable at a local scale, and there are significant variations in exposure and vulnerability across marine ecosystems (e.g., coral reefs versus deep sea). Additionally, the terrestrial human footprint map does not include climate change stressors.

### Analysis

We identify coastal regions as the transition between terrestrial and marine environments based on the 1 km^2^ resolution pressure maps, and represented as points at approximately 1 km distance intervals. We defined a 50 km radius buffer around each point which, following Fang et al. 2018, captures important processes that occur in the coastal zone, including tidal, breeding, and foraging migration of neritic animals, stranded dead marine products on the shore, bird moving foraging, river nutrient transport, and saltwater intrusion (Appendix S1). Spatial error in the location of the coastline points is small relative to the radius within which pressures are quantified.

Intact coastal regions were identified by quantifying the proportion of intact land and sea areas within the 50 km radius of each coastal point (pixel in the human footprint dataset). Any location containing less than 500 cells of either land or sea within the 50 km radius circle (and 30 km radius, 5 cells for the 10 km radius – see Sensitivity Analysis) was omitted from the analysis as the estimate of the proportion of intact area may be unreliable. This occasionally arises as a result of inconsistencies in the mapping of coastlines between the terrestrial and marine data sources. The average of the land and sea intact area proportions was used to characterise the intactness of land sea connected pixels. We divided this final combined metric into five equal bins (0-0.2, 0.2-0.4, 0.4-0.6, 0.6-0.8 and 0.8-1) for reporting purposes. Using the global coastal intactness estimates, we summarise the distribution of coastal intactness by nation and for key coastal ecosystems, and assess their global protection status (Appendix S2).

### Units of analysis

We assessed the intactness of coastal regions in proximity to tidal flats (Murray et al. 2019), saltmarshes (Mcowen et al. 2017), mangroves (Bunting et al. 2018), seagrasses (UNEP-WCMC 2003), estuaries (Alder 2003), kelp forests (Mora-Soto et al. 2020), coral reefs (UNEP-WCMC et al. 2018), savannah (Jung et al. 2020), deserts (Jung et al. 2020), rocky areas (Jung et al. 2020) and forests (Jung et al. 2020). This was achieved by buffering the polygons representing each habitat type by 50 km (a radius equal to the radius used to quantify coastal intactness) and summarising the distribution of coastal intactness values within the buffer. Hence, this is a measure of the intactness of the coastal regions influencing these systems (Ray 1991; Fang et al. 2018), not of intactness within or around each of these habitat types. To delineate national borders we used GADM national boundaries (Global Administrative Areas 2012).

Data on protected area location, and boundary of protected areas were obtained from the June 2019 version of the World Database on Protected Areas (WDPA; UNESCO 2020). We incorporated into the June 2019 version of WDPA 768 protected areas (1,425,770 km^2^) in China (sites that were available in the June 2017 version of WDPA, but not publicly available thereafter). Following the WDPA best practice guidelines (www.protectedplanet.net/c/calculating-protected-area-coverage) and other global studies (Maxwell et al. 2020), we included in our analysis only protected areas from the WDPA database that have a status of ‘Designated’, ‘Inscribed’ or ‘Established’, and removed all points and polygons with a status of ‘Proposed’ or ‘Not Reported’. We also removed all points and polygons designated as ‘UNESCO MAB Biosphere Reserves’, as these do not meet the IUCN definition of a protected area. We buffered the point feature class in accordance to the point’s area as stated in the ‘REP_AREA’ field, and merged the buffered points with polygons to create one polygon layer. To reduce computational burden, we removed redundant vertices (tolerance was set at 1000 m) in the polygon layer.

### Sensitivity analysis

We carried out a sensitivity analysis in relation to our definition of the coastal region, with buffer sizes of 30 km and 10 km, rather than 50 km. We found broadly similar patterns in the distribution and relative frequency of intactness categories among these radii (Appendix S3 and Appendix S5).

For the terrestrial human footprint, an ecologically relevant threshold to define intact habitat has been previously established (Di Marco et al. 2018). However, in the marine realm, a threshold is yet to be ecologically defined and validated. We therefore also carried out our analysis over a range of thresholds and definitions of intact in the marine environment. In the main manuscript we present results where the threshold is set to below the 40% quantile. Our sensitivity analysis included any value below the 20% quantile (excluding climate change stressors), the average of all stressors for the year 2013 for below the 20% and 40% quantile, the average of all stressors for the year 2013 for below the 20% and 40% quantile but excluding climate change stressors, and the full cumulative human impact map for the year 2013 (which includes climate change stressors) for below the 20^th^ and 40^th^ quantile. We found similar trends between the five intactness categories across all definitions of intact in the marine environment. Climate change stressors are the predominate driver of human pressure in the marine environment, their inclusion quite obviously shifted the positioning of the 40^th^ quantile compared to when they were excluded. This changed the results slightly, for example, this changed the percentage of coastal regions within the 0.8-1 intactness category from 15.5% (when climate change stressors were included) to 9.04%. See Appendix S4, and Appendix S5 for results.

## Results

### The intactness of Earth’s coastal regions

Using this assessment approach we find that no coastal region is free from human influence (i.e., 100% intact) and only 15.5% of all coastal regions can be considered “low” in anthropogenic pressure (80 – 100% intact; Fig. 1). Conversely 14.0% is exposed to extreme human pressure (0% intact), and 47.9% of coastal regions are exposed to high human pressure 0-20% intact. These coastal regions with high levels of human pressure are located across Earth, but are more concentrated in tropical and temperate regions (Fig. 1). There are more coastal regions that have low intactness in the marine realm but relatively high intactness in the terrestrial realm, than high intactness in both realms (Fig. 2).

**Figure 1.**
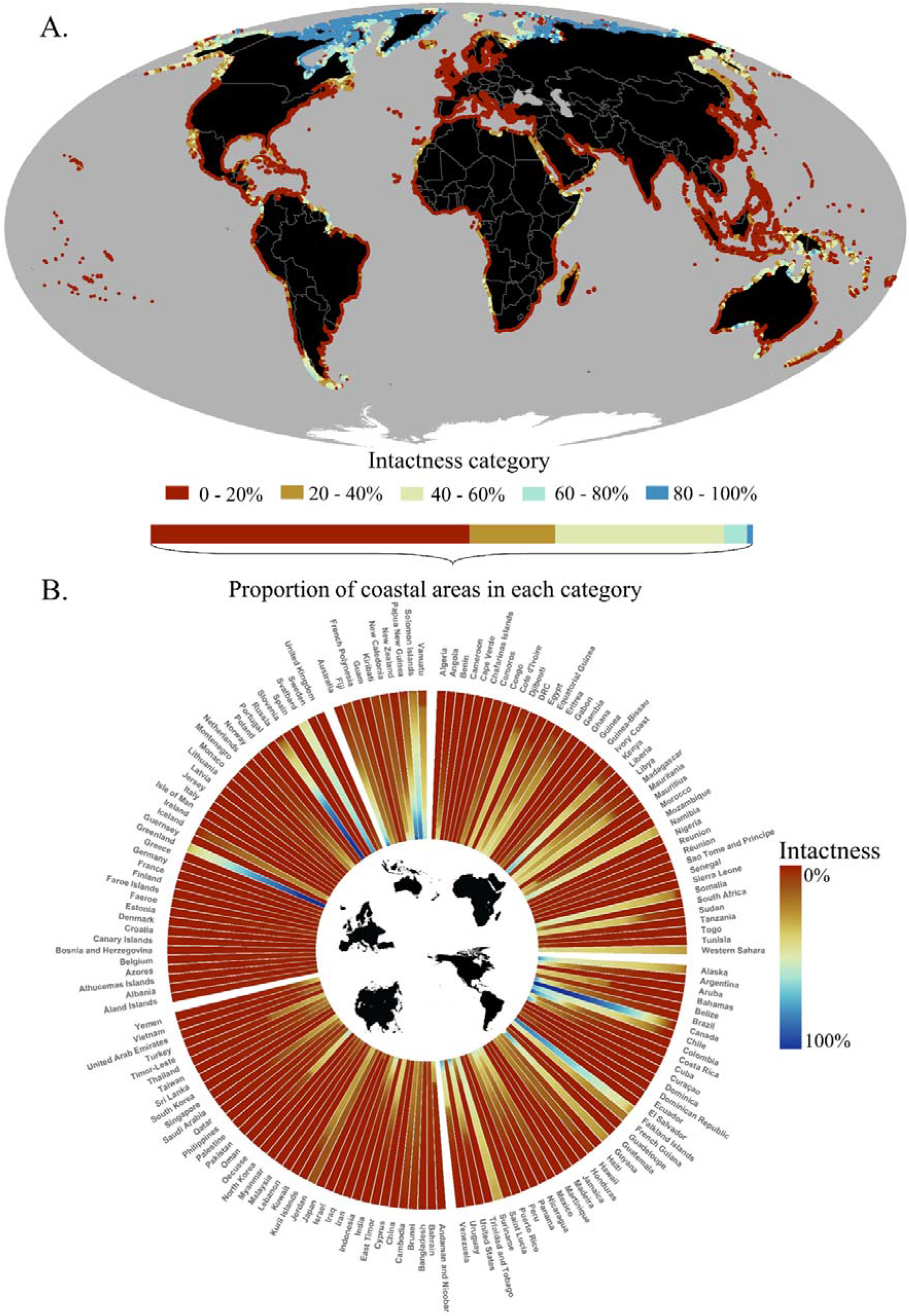
Intactness of Earth’s coastal regions (A) and the proportion of each country’s coastal regions that are intact (across a scale of 0-100%; B). For the terrestrial realm we define intactness using the terrestrial human footprint (under a threshold of <4, representing a reasonable approximation of when anthropogenic land conversion has occurred to an extent that the land can be considered human-dominated; Williams et al. 2020), and for the marine realm we use the cumulative human impact dataset (under a threshold of the 40% quantile – and excluding climate change pressures; Halpern et al. 2019).

**Figure 2.**
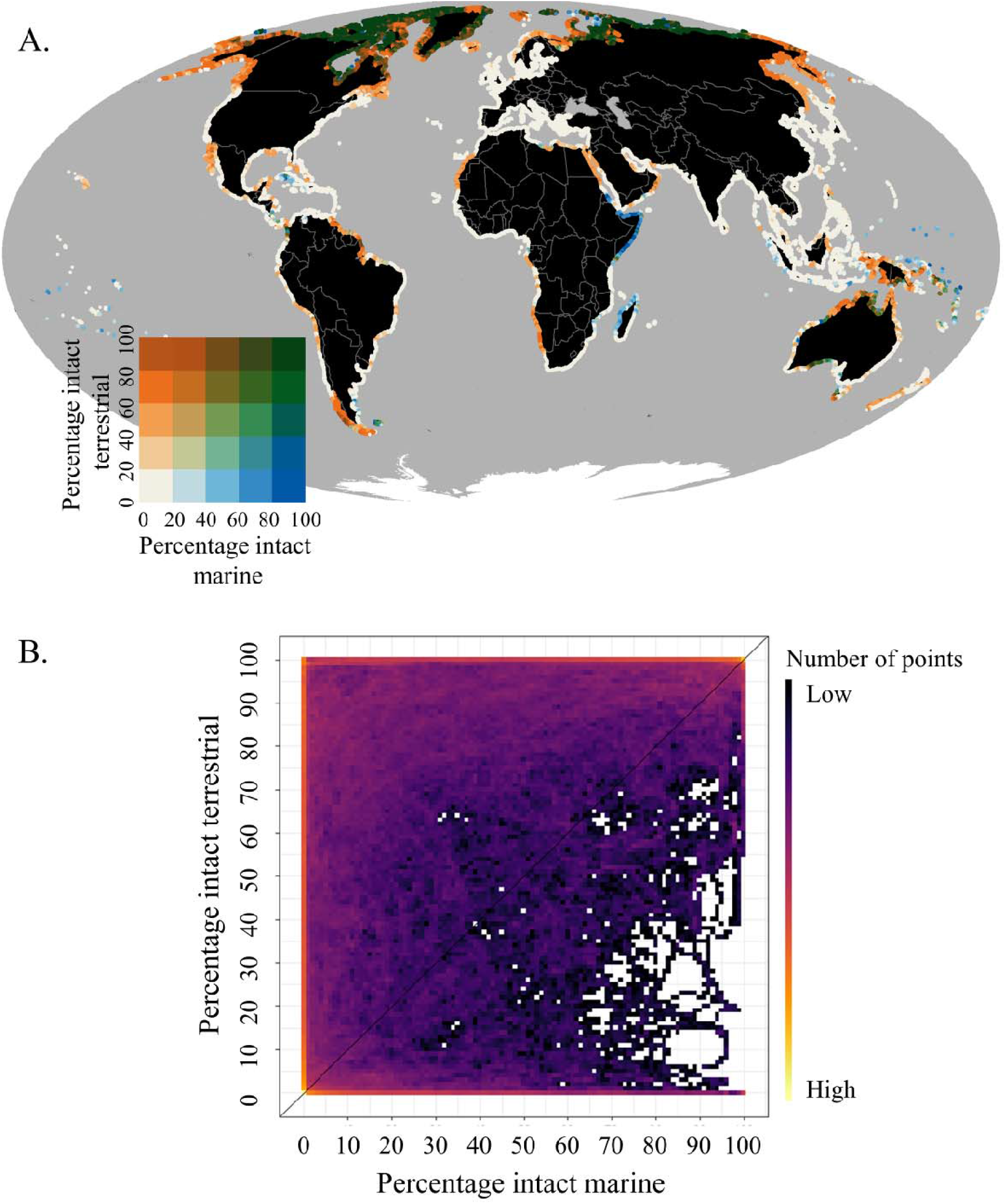
(A) The spatial relationship between intactness in the terrestrial and the marine realms. (B) A heat map of the number of points (ranging from 1 to 249,136) and the corresponding proportion within the 50 km buffer that is intact in the marine realm (x-axis; under a threshold of the 40% quantile (Halpern et al. 2019)) and in the terrestrial realm (y-axis; under a threshold of <4 (Williams et al. 2020)).

Almost all of the most intact coastal regions are located in Canada, Russia, and Greenland (Fig. 1B). Canada is responsible for the largest expanse of coastal region that remains under very low anthropogenic pressure, with 53.4% of its coastal regions (>60,855 km or 7.93% of all coastal regions) falling within the highest intactness category (>80% intact), followed by Russia 40.7% (>34,737 km or 4.52%), and Greenland 44.1% (>19,176 km or 2.50%; Fig. 1B). Their relatively intact condition can likely be attributed to their remoteness from major urban and industrial centres, and inaccessibility during winter months (Halpern et al. 2008). We found 12 nations contain coastal regions that remain >80% intact, and a further 9 contain coastal regions that remain relatively intact (60-80% intact), with large expanses located in Chile, Australia, the United States, Svalbard, Indonesia, Papua New Guinea, the Falkland Islands, the Solomon Islands, and Brazil. At the other extreme, we found that all coastal regions of 26 nations are highly exposed to human pressures (i.e., 0% intact). Many of these were island nations including Singapore, Dominica, and Aruba, but also included mainland nations in Africa, and Asia (Fig. 1B; Appendix S7).

### The intactness of coastal ecosystems

We quantified human pressure across 11 coastal ecosystems (forests, rocky areas, savannah, desert, coral reefs, estuaries, kelp forests, mangroves, salt marshes, seagrasses, tidal flats) and found that more than 60.1% of the coastal regions containing these ecosystems is under high levels of human pressure (0-20% intact; Fig. 3). Human pressure is highest across the coastal regions with seagrasses, savannah, and coral reefs; 80.0% of coastal regions adjacent to seagrass (>195,775 km), 77.3% adjacent to savannah (>138,594 km), and 73.8% adjacent to coral reefs (>172,502 km) are exposed to high human pressures (0-20% intact; Fig. 3). Deserts, forests, and salt marshes are the coastal habitats that have the most area within intact regions (80-100% intact), but this is only 3.90%, 1.21%, and 0.48% of Earth’s coastal regions, respectively (Fig. 3; Appendix S6).

**Figure 3.**
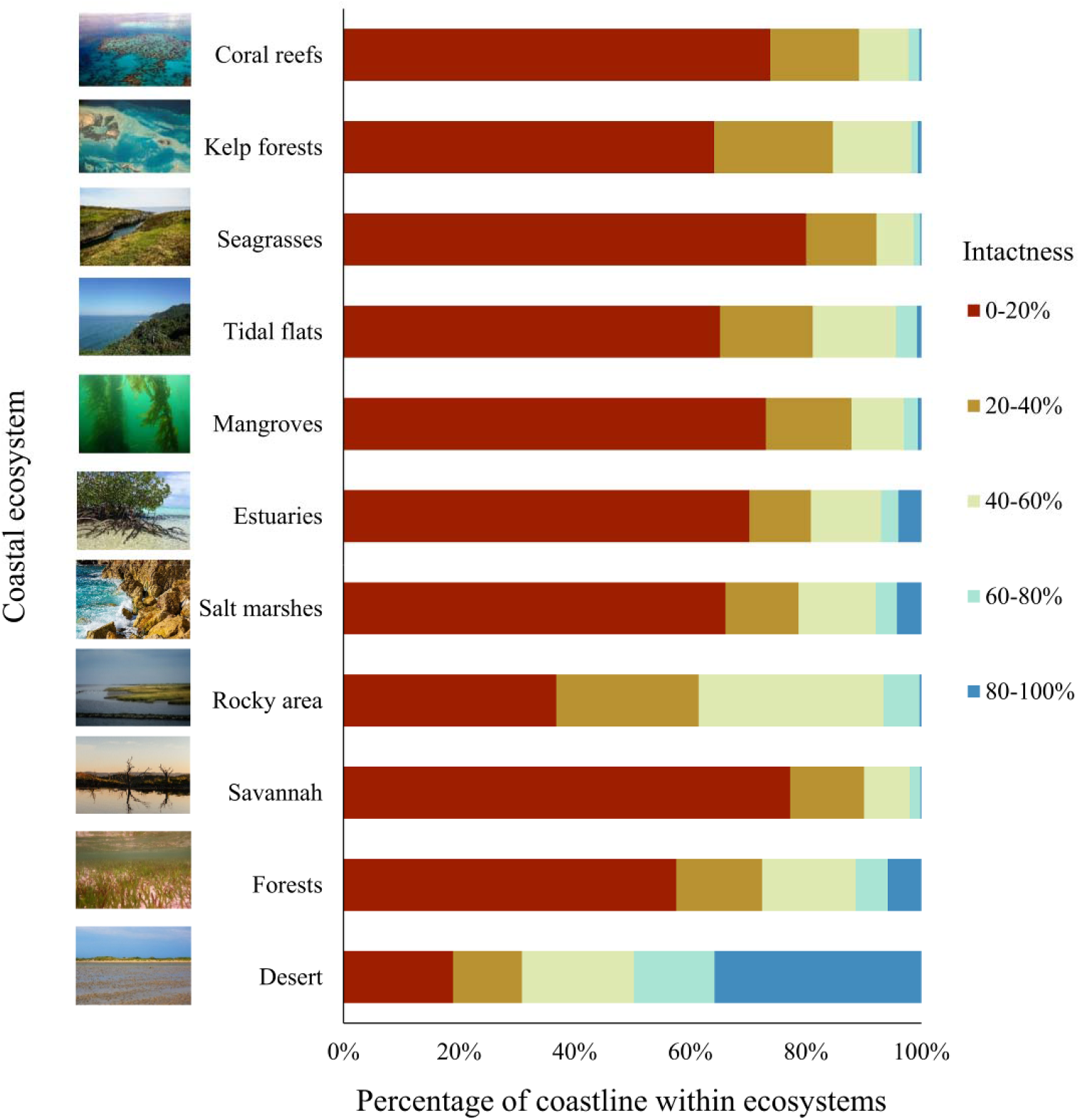
Distribution of intactness values of coastal regions in proximity (< 50km) to eleven coastal ecosystem types. Photo credits - Coastal forest © Brooke Williams, Kelp forest and Seagrasses © Megan Saunders, Mangroves © Leslie Roberson, all other images are © creative commons.

### Current levels of coastal protection

Of the 16.4% of coastal regions falling within designated, inscribed, or established (World Database on Protected Areas; UNESCO 2020) protected areas (either in the terrestrial or marine environment), 43.3% are exposed to high human pressures (0-20% intact). The United States, Russia, Canada, and Greenland, have the most coastal region protected (in terms of area), whereas Greenland, Canada, and Svalbard have the most coastal regions of high intactness protected (Fig. 4). We found 61.9% of protected coastal regions (> 257,004 km) was protected in both the terrestrial and marine environments, the remaining 38.1% was protected only in one realm (11.2% marine, 88.8% terrestrial).

**Figure 4.**
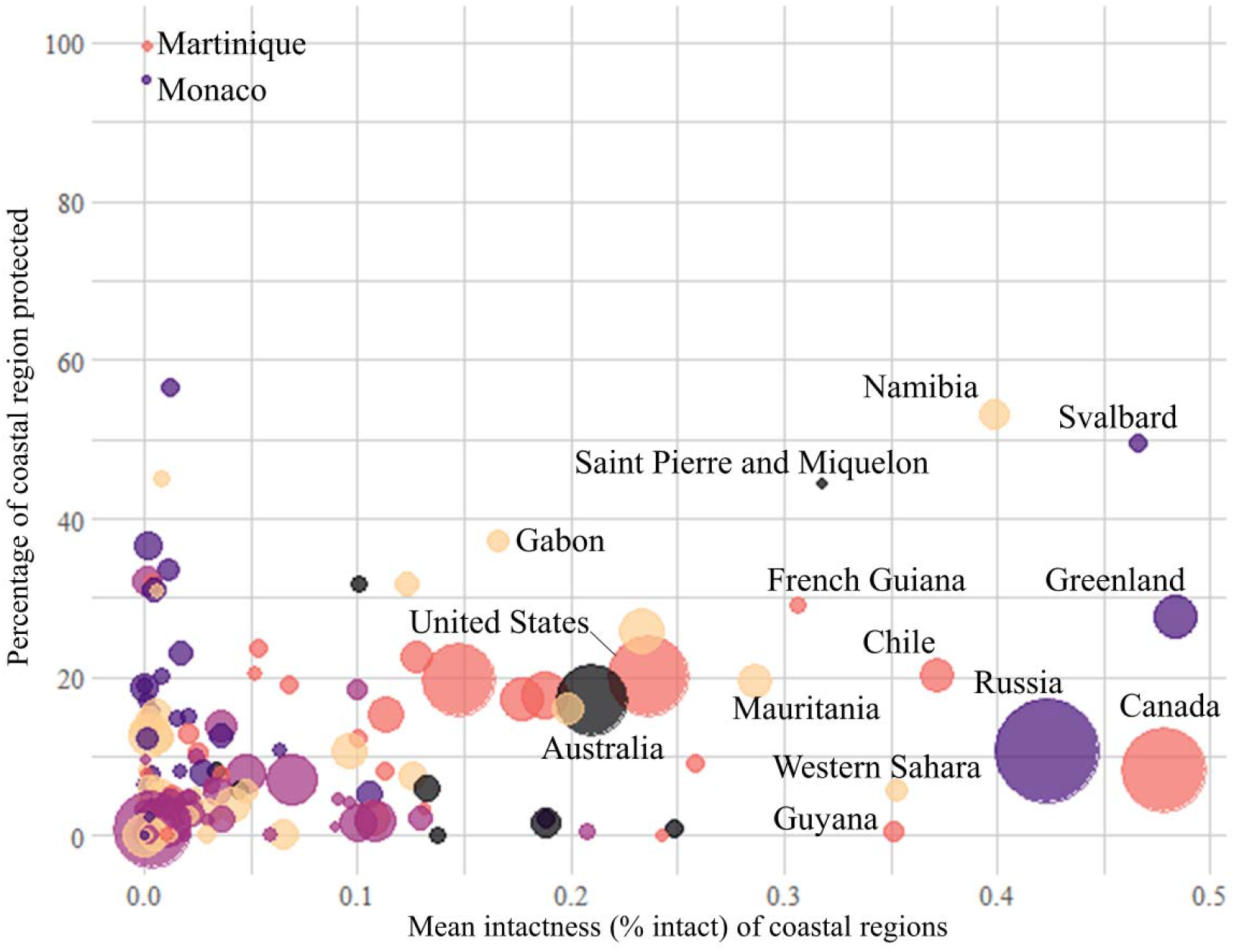
The percentage of each countries coastal region that is protected (World Database on Protected Areas (WDPA; UNESCO 2020) that have a status of ‘Designated’, ‘Inscribed’ or ‘Established’) plotted against the mean intactness of coastal regions. Bubble sizes represent relative country size and suites of colours represent bio-geographic regions (Asia – light purple, Oceania - black, Europe – dark purple, America – pink, or Africa - cream).

## Discussion

It is safe to say intact coastal regions are now rare. This has profound implications for coastal biodiversity (Hazlitt et al. 2010; Rogers & Mumby 2019), and for humanity, as we rely on functioning coastal ecosystems for ecosystem services such as climate change mitigation, food provision, and storm protection (Mcleod et al. 2011; Pendleton et al. 2012; Ferrol-Schulte et al. 2013). Many of the coastal regions that remain intact are at higher latitudes, so broad-scale restoration is required across much of Earth’s coastal regions. However, where and how to restore, protect, or manage varies depending on the levels of human pressure coastal regions are experiencing (Darling et al. 2019).

Relatively intact coastal regions will require different conservation implementation strategies to conserving the last remaining intact pockets (areas of low human pressure surrounded by areas of higher human pressure), which can provide benefits to surrounding locations of lower integrity (Cinner et al. 2020). For example, in northwest Greenland conserving long stretches of sea ice, arctic water, and glacier habitats will require enhancing environmental governance and laws around encroaching development and addressing climate change (climate change stressors are not included in the results of the main manuscript; however, we include them for the marine realm in the sensitivity analysis (see Appendix S4 and Appendix S5)) (Nuttall 2020). Here, and across many other coastal regions, strengthening Indigenous peoples’ involvement in managing coastal environments will be vital to long-term coastal sustainability. In contrast, conserving the remaining intact pockets of coastlines, like those on the coastal region encompassing Collingwood Bay in Papua New Guinea, which is relied on by local communities for ecosystem services (Poloczanska et al. 2011), and areas in the Tambelan Archipelago in Indonesia, where the coastal ecosystems are known nurseries for fish species (Yonvitner & Fahmi 2012), will rely on specific management actions by local communities. In Collingwood Bay land owners have been battling illegal logging for decades - here conservation success will take the form of complex socio-political action to address largely land-based stressors (McDonnell et al. 2017). Efforts such as the analysis presented here can help differentiate this spectrum of human pressure at a broad scale, to drive localized assessments to inform actions on the ground (Cross et al. 2012).

Encouragingly we found that 61.9% of protected coastal regions are protected in both the terrestrial and marine realms, rather than protection occurring in just one realm. Examples include the Patagonia Fjords of southern Chile, some locations along the Australian coast (including many wetland ecosystems) adjacent to the Great Barrier Reef, and the Iguape-Cananéia-Paranaguá estuary in the state of Paraná in Brazil. Management strategies here consider the coastal interface through community and local engagement to better understand degrading processes (Anbleyth-Evans et al. 2020), retention of riparian coastal ecosystems to prevent nutrient and sediment run-off from surrounding agricultural lands (Kroon et al. 2016), and effectively managing fishing resources (Mendonça et al. 2010). These examples should be showcased and implemented more broadly.

Protected areas alone cannot mitigate all threats and are not enough to safeguard Earth’s coastal regions. As the 16.4% of coastal regions under formal protection are under high levels of human pressure, increasing well-resourced protected areas is an important priority (Fraschetti et al. 2009), but they must be accompanied by other effective area-based conservation measures (OECMs) (Cinner et al. 2020), and non-area based management (Wenger et al. 2018a) to deliver biodiversity and ecosystem service benefits. OECMs are likely going to be an increasingly important coastal management strategy, as they can be an opportunity to conserve nature while ensuring community rights are recognised and that communities are enfranchised to manage their own resources while delivering biodiversity and ecosystem service benefits (Dudley et al. 2018). This is particularly acute in the intact coastal areas of the arctic, which are the homelands for diverse groups of indigenous peoples, each with their own distinct cultures, histories and livelihood practices such as reindeer herding, subsistence whale and seal hunting, and commercial fisheries. In these cases, it will be crucial to work with these communities to maintain the ecological integrity of these intact coasts in the face of industrial development pressures, while harnessing their traditional ecological knowledge for the process of adapting to climate change (Fondahl et al. 2015; Gassiy & Potravny 2019). Non-area based management approaches such as mitigating land-use change to prevent increased pollution run-off (Hamilton et al. 2017), or enhanced regulation of degrading activities (Wenger et al. 2018b) will also play a crucial role. These conservation actions must be implemented with careful consideration of the often over looked land-sea connection that coastal regions encompass, rather than independent land or sea initiatives (Jupiter et al. 2017).

Given the widespread human pressure we have revealed across coastal regions (Fig. 1, Fig. 3), there is a fundamental role for active restoration in many nations (Saunders et al. 2017). Priorities for coastal restoration should be informed by levels of human pressure, as it will depend on the removal of threats (Borja et al. 2010) and targeted management of anthropogenic activities such as nutrient run-off (Duarte et al. 2020). Sea-level rise as a consequence of climate change is also leading to coastal flood and erosion risks, inciting efforts to restore defence ecosystems either to their natural state or replicate their function through artificial structures (Pontee et al. 2016). But these restoration actions must also be informed by regional assessments of ecological integrity and feasibility of success (Bayraktarov et al. 2016). Some locations may be so degraded that the cost-benefit ratio of active restoration may be high, and other types of conservation actions such as passive restoration, off-setting or relocation of species may be warranted (Gayle et al. 2005).

Our analysis is the first to integrate the terrestrial and marine human pressure maps for coastal regions. As both maps were created independently, there are inherent limitations in their congruence. As identifying intact areas requires finding those areas that have little to no impact across all human activities (Jones et al. 2018), for the marine realm we regard intact as any value below the 40% quantile in the cumulative human impact map. Future studies may follow assessments that have been carried out on land and empirically assess the ecological significance of this threshold (Di Marco et al. 2018). In addition, climate change stressors, including ocean acidification, sea surface temperature, and sea-level rise, were omitted from the marine cumulative human impact dataset for the purpose of this analysis (but see the sensitivity analysis, Appendix S4 and Appendix S5 for results when they are included). However, climate change threatens most coastal regions through changes to biophysical and socioeconomic processes that can be difficult to predict. This analysis therefore represents an optimistic assessment of ecosystem intactness in the context of future climate change impacts that are likely to arise with increasing frequency over the coming decades (IPPC 2018).

Our research shows that humanity’s impact on Earth’s coastal regions is severe and widespread. In order to meet global conservation and sustainability goals, it is crucial that nations implement conservation activities to retain their remaining intact coastal regions. Nations like Russia, Canada and Greenland can play a significant role by proactively protecting the last great intact coastal regions on Earth. Most coastal nations have small pockets of intact coastal regions and it is particularly important to maintain these environments. Our research also shows that it is now critical the global community set specific restoration targets for coastal regions, which would support the United Nation’s Decade for Ecosystem Restoration efforts that are underway.

## Supporting Information

Definition of coastal region (Appendix S1), a flow diagram of the methods used to calculate coastal region intactness (Appendix S2), coastal region intactness using a buffer size of a) 10 km and b) 30 km as opposed to 50 km as a sensitivity analysis (Appendix S3), coastal region intactness using a range of thresholds and definitions of intact in the marine environment as a sensitivity analysis (Appendix S4), coastal region intactness (number of points) using a range of buffers sizes to define coastal region, and definitions of intact in the marine environment from the sensitivity analysis to determine percentage intact (Appendix S5), distribution of intactness values (number of points) of coastal regions in proximity (< 50 km) to 11 coastal ecosystem types (Appendix S6), and the amount (number of points) of each country’s coastal regions and their percentage intactness (0-100%) (Appendix S7). The authors are solely responsible for the content and functionality of these materials. Queries (other than absence of the material) should be directed to the corresponding author.

## Acknowledgements

BAW and LAR were supported by an Australian Government Research Training Program Scholarship. CJK was funded by a University of Queensland Fellowship and the Australian Research Council. The authors declare no competing interests.

## Notes

### Competing Interest Statement

The authors have declared no competing interest.

